# Intercellular Arc signaling regulates vasodilation

**DOI:** 10.1101/2020.08.13.250209

**Authors:** Paulino Barragan-Iglesias, June Bryan de la Peña, Tzu-Fang Lou, Sarah Loerch, Nikesh Kunder, Tarjani Shukla, Lokesh Basavarajappa, Jane Song, Salim Megat, Jamie K. Moy, Andi Wanghzou, Pradipta R. Ray, Jason Shepherd, Kenneth Hoyt, Oswald Steward, Theodore J. Price, Zachary T. Campbell

**Affiliations:** University of Texas at Dallas, School of Behavioral and Brain Sciences, Richardson, TX, USA; Department of Physiology and Pharmacology, Center for Basic Sciences, Autonomous University of Aguascalientes, Aguascalientes, Mexico; University of Texas at Dallas, Department of Biological Sciences, Richardson, TX, USA; University of Texas at Dallas, Department of Bioengineering, Richardson, TX, USA; Janelia Research Campus, Howard Hughes Medical Institute, Ashburn, VA, USA; Department of Neurobiology and Anatomy, University of Utah, Salt Lake City, UT, USA; Reeve-Irvine Research Center, Departments of Anatomy & Neurobiology, Neurobiology & Behavior, Neurosurgery, School of Medicine, University of California Irvine, Irvine, CA, USA; Center for Advanced Pain Studies, University of Texas at Dallas, Richardson, TX, USA

## Abstract

Injury responses require communication between different cell types in the skin. Sensory neurons contribute to inflammation and can secrete signaling molecules that affect non-neuronal cells. Despite the pervasive role of translational regulation in nociception, the contribution of activity-dependent protein synthesis to inflammation is not well understood. To address this problem, we examined the landscape of nascent translation in DRG neurons treated with inflammatory mediators using ribosome profiling. We identified the activity-dependent gene, Arc, as a target of privileged translation *in vitro* and *in vivo*. Inflammatory cues promote local translation of Arc in the skin. Arc deficient mice display exaggerated paw temperatures and vasodilation in response to an inflammatory challenge. Since Arc has recently been shown to be released from neurons in extracellular vesicles, we hypothesized that intercellular Arc signaling regulates the inflammatory response in skin. We found that the excessive thermal responses and vasodilation observed in Arc defective mice are rescued by injection of Arc-containing extracellular vesicles into the skin. Our findings suggest that activity-dependent production of Arc in afferent fibers regulates neurogenic inflammation through intercellular signaling.

**HIGHLIGHTS:**

- Ribosome profiling identifies Arc as a target of activity-dependent translation
- Arc is present in the DRG, spinal cord, and skin
- Induced Arc biosynthesis in skin requires the presence of afferent fibers
- Arc-deficient mice have exaggerated inflammation that is rescued by Arc-containing extracellular vesicles

## INTRODUCTION

The skin forms a protective barrier to pathogens and is an essential point of contact between vertebrates and their environment. Skin is the single largest sensory organ due to innervation by nerve fibers that detect a broad range of stimuli. Sensory fibers communicate information from the periphery to the central nervous system. These cues can illicit behavioral responses that promote injury avoidance. A second critical function enabled by sensory neurons is regulated release of signaling molecules that influence blood flow, cellular proliferation, and immune response (Dubin and Patapoutian, 2010; Pinho-Ribeiro et al., 2017; Toda et al., 2008). While a handful of the factors released by afferent fibers have been identified, the mechanisms that govern their biosynthesis are largely unknown.

Afferent fibers play a central role in neurogenic inflammation. Neuropeptides, including calcitonin gene related protein (CGRP) and substance P, are released from afferent fibers (Brain and Williams, 1989; Saria, 1984). While both are potent vasodilators, their mechanisms of action differ. While CGRP promotes vasodilation, substance P increases capillary permeability. The impact of communication between nociceptors and non-neuronal cell types on injury repair is profound. Nociceptive mediators stimulate recruitment of immune cells and contribute to both innate and adaptive immunity (Pinho-Ribeiro et al., 2017). Regulated protein synthesis is critical in sensory neurons. Translational inhibition has been shown to reduce pain associated with inflammation, injury, neuropathy, and migraine (Avona et al., 2019; Barragan-Iglesias et al., 2019; Ferrari et al., 2013a, 2013b, 2015; Megat et al., 2019; Moy et al., 2017). It is unclear which transcripts are subject to activity-dependent translation in afferent fibers. In the central nervous system, neuronal activity induces the expression of a number of effectors collectively referred to as Immediate Early Genes (IEG) (Guzowski et al., 1999; Link et al., 1995; Lyford et al., 1995; Rosen et al., 1998; Vann et al., 2000).

To understand the impact of inflammatory mediators on the landscape of nascent translation in afferent neurons, we used ribosome profiling to analyze cultured dorsal root ganglion (DRG) neurons (Ingolia, 2010). Application of the inflammatory mediators Nerve Growth Factor (NGF) and Interleukin 6 (IL-6) induces persistent changes in the activity of DRG neurons (Chang et al., 2016; De Jongh et al., 2003; Obreja et al., 2018; Svensson, 2010). Based on altered patterns of ribosome occupancy, we identify numerous mRNAs with altered rates of translation. Intriguingly, we found a robust increase in biosynthesis of the prototypical IEG, Arc. Arc is critical for long-term forms of synaptic plasticity and memory (Shepherd and Bear, 2010; Plath et al., 2006). Moreover, dendritic translation of Arc mRNA is tightly correlated with activation of circuits implicated in learning and memory (McIntyre et al., 2005). A surprising connection has recently been established between Arc and retroviruses (Erlendsson et al., 2020; Pastuzyn et al., 2018). Arc protein is homologous to the HIV Gag protein and forms virus-like capsids. Arc mediates intercellular mRNA transport between neurons and a range of recipient cell types through Extracellular Vesicles (EVs). In *Drosophila*, the Arc homologue dArc1 transits from presynaptic neuromuscular synapses to muscle (Ashley et al., 2018). However, a precise physiologic function for intercellular communication mediated by Arc capsids remains to be determined in mammals. Here, we show that inflammatory signals promote rapid translation of Arc *in vivo* and *in vitro*. Mice that lack Arc have exaggerated thermal responses and vasodilation following an inflammatory challenge in the skin. Both are rescued by injection of Arc-containing EVs. Our data suggest a new role for Arc as a mediator of intercellular signaling in peripheral neuroinflammation.

## RESULTS

### Ribosome profiling identifies Arc as a target of privileged translation

To profile translation in the DRG, tissues were harvested from cervical (C1) to lumbar (L6) segments of the spinal column and maintained for five days *in vitro* (Figure 1A). Cultures were subjected to either a vehicle treatment or exposed to two inflammatory mediators: NGF and IL-6. These molecules were selected because both rapidly induce nascent protein synthesis and lead to persistent changes in the activity of sensory neurons (Melemedjian et al., 2010). After 20 minutes, the elongation inhibitor emetine was added to each culture to stop translation and cells were lysed (**STAR★Methods**). These samples were used to generate libraries for two types of sequencing. As a control for changes in mRNA abundance that could reflect transcription responses or changes in stability, mRNA abundance was quantified with high-throughput sequencing. To generate these libraries, ribosome RNAs were depleted and 3’ end sequencing was conducted using Quantseq (Moll et al., 2014). The majority of the sample was used to generate the other type of library consisting of ribosome protected footprints. After translation was arrested with emetine, lysates were nuclease treated, small RNAs were purified, and adapters were added in a ligation independent manner (Hornstein et al., 2016). This process was repeated for a total of five independent replicates. The high-throughput sequencing experiments were highly reproducible both for ribosome profiling and for RNA-seq (Pearson’s R > 0.98, Figure 1B-C). We found that the majority of ribosome protected fragments mapped to annotated coding regions (Figure 1D). The footprints displayed a triplet periodicity consistent with ribosomal translocation (Figure S1A).

**Figure 1.**
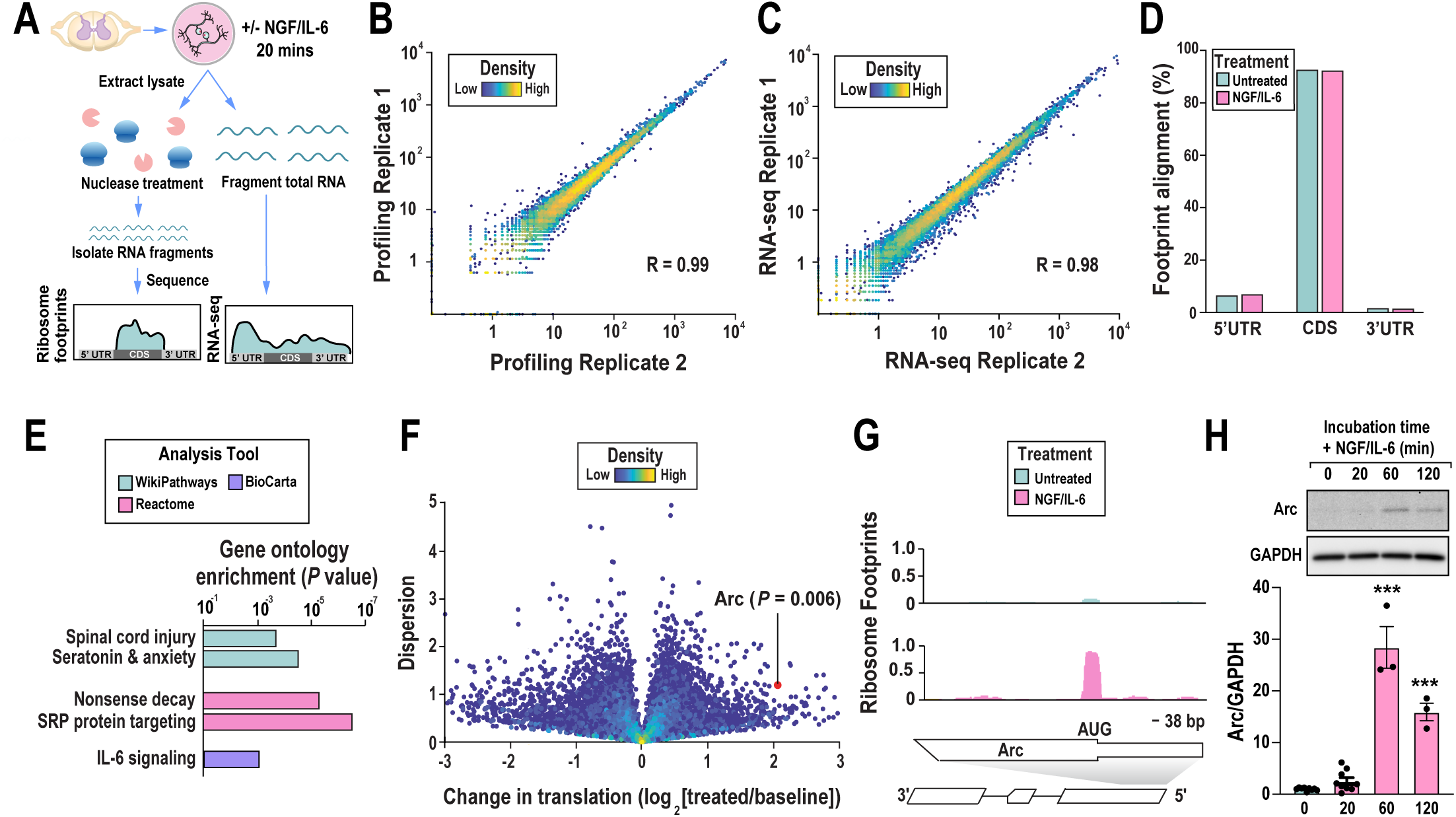
Ribosome profiling identifies Arc as a rapidly-translated gene. (A) Schematic representation of the ribosome profiling approach applied to DRG neurons. (B) Intra-experimental correlations for ribosome profiling in the inflammatory mediator treatment group. (C) Intra-experimental correlations for RNA-Seq in the inflammatory mediator treatment group. R values correspond to Pearson’s correlations in B-C. (D) Aggregate sites of ribosome protected footprints in 5’ untranslated regions, coding sequences, and 3’ untranslated regions. (E) Gene ontology analysis of genes with significant translational enhancement was conducted with Enrichr (Kuleshov et al., 2016). (F) A volcano plot depicting changes in translation (log_2_ ≥ –1.5; false discovery rate < 0.05) across the transcriptome. The ratio of ribosome density following treatment with the inflammatory mediators divided by the vehicle levels is plotted against sample dispersion. Gene density is shaded according to the inset bar. P-values for Arc were determined using a two-tailed Student’s t-test. (G) Traces of the 5’ end of *Arc*. Scales are indicated with bars. (H) Application of NGF/IL-6 to primary culture DRG neurons up-regulated Arc protein in a time course of 0-120 min. n = 9 for 0- and 20-min groups. n = 3 for 60- and 120-min groups. Ordinary one-way ANOVA: F_(3, 20)_ = 95.79, P < 0.0001. Dunnett’s multiple comparisons test: 0 vs 20 min, ***P < 0.001; 0 vs 120 min, ***P < 0.001.

To determine if the treatment induced larger changes in mRNA abundance or translation, we compared differences in each dataset. We found that translation of 217 mRNAs was increased by more than 1.5-fold (Figures 1F, S1B, Tables S1 and S2). In contrast, fewer than 100 mRNAs were increased in abundance (Figure S1B). Therefore, we focused our analysis on targets of differential translation. To determine if the preferentially translated mRNAs encoded products with related functions, we performed an ontological analysis. We found enrichment for several processes including: spinal cord injury (Figure 1E, P = 0.002), IL-6 signaling (P = 0.009), and serotonin and anxiety-related events (P = 0.0003) (Kuleshov et al., 2016). To determine what features were commonly observed in this set of transcripts we examined 5’ UTR length and GC content (Figures S1C-D). While we did not detect an overt difference, we found a nine-base motif that was significantly enriched in the 5’ UTR (Fisher’s exact test, P = 0.02, Figures S1E-F). This sequence motif was reminiscent of an mTOR responsive element termed the pyrimidine-rich translational element (Hsieh et al., 2012). Based on ribosome profiling, we conclude that cultured DRG neurons preferentially translate genes linked to neuronal function and memory and are enriched for a specific motif found in the 5’UTR.

Among the transcripts with significant increases in translation was the immediate early gene Arc (Figure 1G). While Arc is expressed in the spinal cord, it is unclear if it is present in DRG neurons (Bojovic et al., 2015; Hossaini et al., 2010; Minatohara et al., 2015; Plath et al., 2006). To validate if Arc is translated in response to inflammatory mediators, we conducted immunoblots on cultured DRG exposed to NG/IL-6 (Figure 1H). We found little to no detectable Arc protein prior to addition of NGF and IL-6. However, one hour after addition, Arc protein levels were increased by more than 15-fold. These results suggest that Arc is translated in response to inflammatory mediators *in vitro*.

### Arc mRNA is expressed in large diameter DRG neurons

Next, we sought to determine if *Arc* mRNA is preferentially expressed in a specific class of sensory neuron *in vivo*. To identify cells that express *Arc*, we analyzed single cell sequencing previously conducted on dissociated DRG neurons (Li et al., 2016) (Figure 2A). We qualitatively compared expression of *Arc* to markers for peptidergic (*Calca/CGRP*), non-peptidergic (*P2×3*) and a light chain neurofilament expressed in large diameter neurons (*NELF*). *Arc* is expressed at low levels in a range of cell types but lacks a clear bias in the single cell sequencing experiments. To examine the expression of *Arc* in tissues, we conducted *in situ* hybridization on L4-L5 DRG tissues harvested from animals exposed to the inflammatory mediators (henceforth referred to as ipsilateral) or negative controls collected from the same animals on the uninjected side of the body (henceforth contralateral) (Figure 2B). Using the same series of markers, we find instances where *Arc* expression coincides with each marker. However, *Arc* levels are highest in large diameter neurons that express NF200 and peptidergic neurons that express *Calca* (Figure 2C). *Arc* expression was found to be increased by strong nociceptive stimuli in the spinal cord (Hossaini et al., 2010). However, we found no difference in *Arc* expression in the DRG or spinal cord (Figure 2D-E) following injection of NGF/IL-6. As a control for the specificity of our *in situ* hybridization probes, we repeated our experiments with a negative control RNA probe for *Arc* and found little to no detectable signal in either the DRG or spinal dorsal horn (Figure S2A-C). Reporter mice have been generated that express a translational fusion between Arc and eGFP (Steward et al., 2017). Arc-GFP is present in DRG neurons and in sciatic nerve fibers in these animals (Unpublished observation).

**Figure 2.**
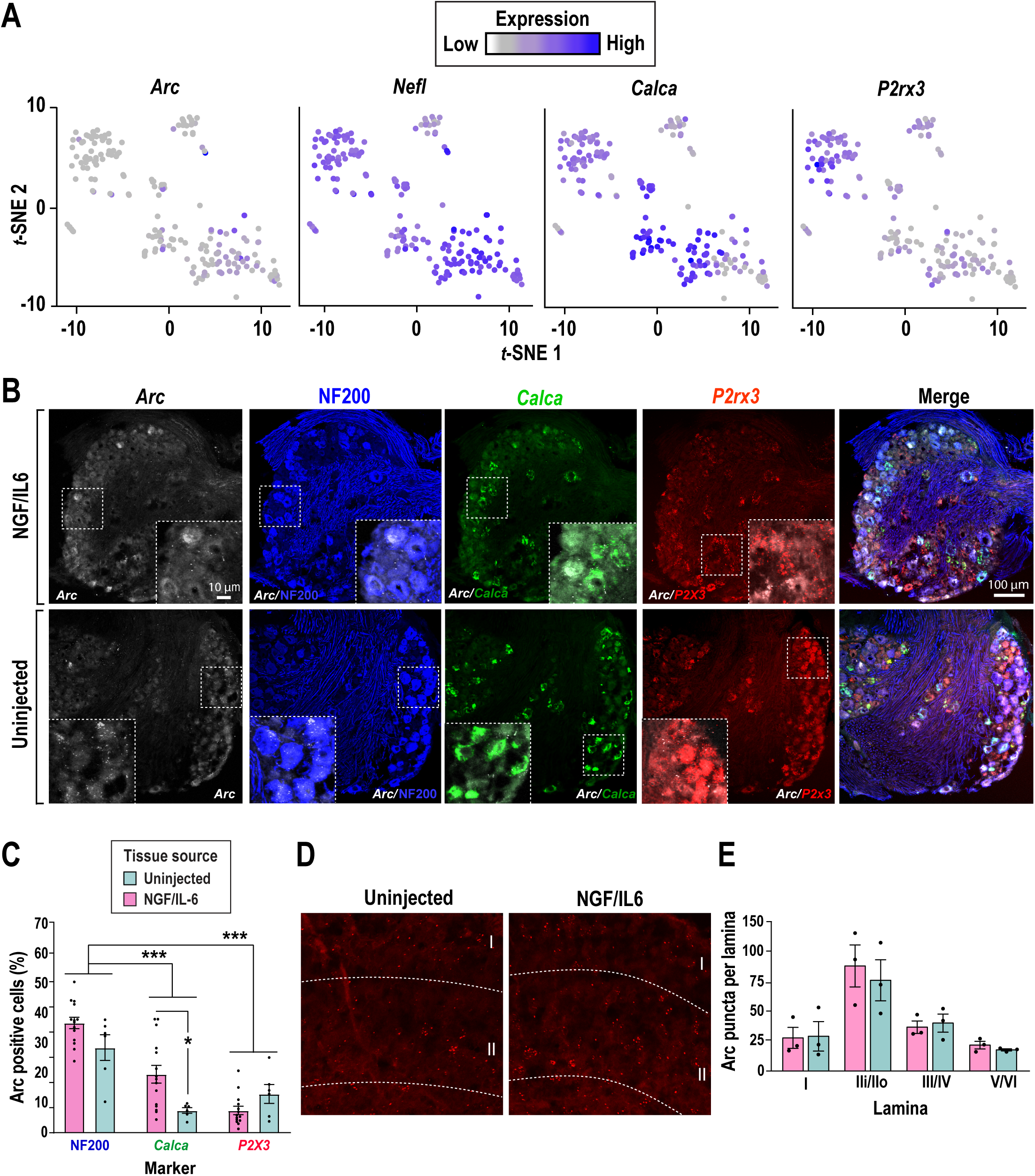
*Arc* is constitutively expressed in peripheral and spinal sites important for nociceptive signaling. (A) Single DRG neuron-sequencing tSNE clusters showing expression of *Arc* with the light neurofilament expressed in large diameter neurons (*Nefl*) and across DRG sensory neurons of peptidergic [*Calca* (CGRP) and non-peptidergic [*P2×3* (P2×3)] sub-clusters (Li et al., 2016). (B) In situ hybridizations of Arc and markers for L4-L5 DRGs together with large diameter neurons expressing neurofilament protein NF200 (NF200, blue), *Calca* (CGRP, green), and neurons expressing *P2×3* (P2×3 red). (C) Quantification of *Arc* expression in the DRG. Expression of *Arc* is greatest in NF200 expressing large diameter neurons. Intraplantar administration of the inflammatory mediators increased *Arc* mRNA in ipsilateral (IPL) DRGs expressing *Calca* (CGRP) mRNA. n=3 animals per group; multiple slices from L4-L5 DRGs. Two-way ANOVA: cell-type factor, F_(2, 55)_ = 35.80, P < 0.0001; treatment factor, F_(1, 55)_ = 4.545, P = 0.0375. Bonferroni’s multiple comparisons test: NF200 vs. CGRP, P < 0.0001; NF200 vs. P2XR, P < 0.0001; CGRP-IPL vs CGRP-CL, *P = 0.0113. (D) In situ hybridization of *Arc* in the dorsal horn of the spinal cord (lamina I-VI). (E) Quantification of D. *Arc* expression is highest in laminal layer II. Intraplantar administration of inflammatory mediators did not significantly impact *arc* mRNA expression.

### Local translation of Arc in the paw

To determine if inflammatory mediators induce rapid translation of endogenous Arc *in vivo*, we examined Arc levels in skin from the hindpaw of injected or uninjected paws one hour after treatment. Immunoblots show that the treatment induces rapid translation of Arc in the ipsilateral paw (Figure 3A). To test if Arc production is the result of *de novo* transcription or translation of pre-existing mRNA, a transcription inhibitor (actinomycin D) was included in the treatment (Figure 3A). Inhibition of transcription did not significantly diminish induced translation of Arc, suggesting that Arc is likely translated from the population of existing mRNA. Next, we asked if Arc translation is induced in either the DRG or the dorsal horn of the spinal cord. In both tissues, Arc is detected, but levels are not increased by NGF/IL-6 treatment (Figures 3B-C).

**Figure 3.**
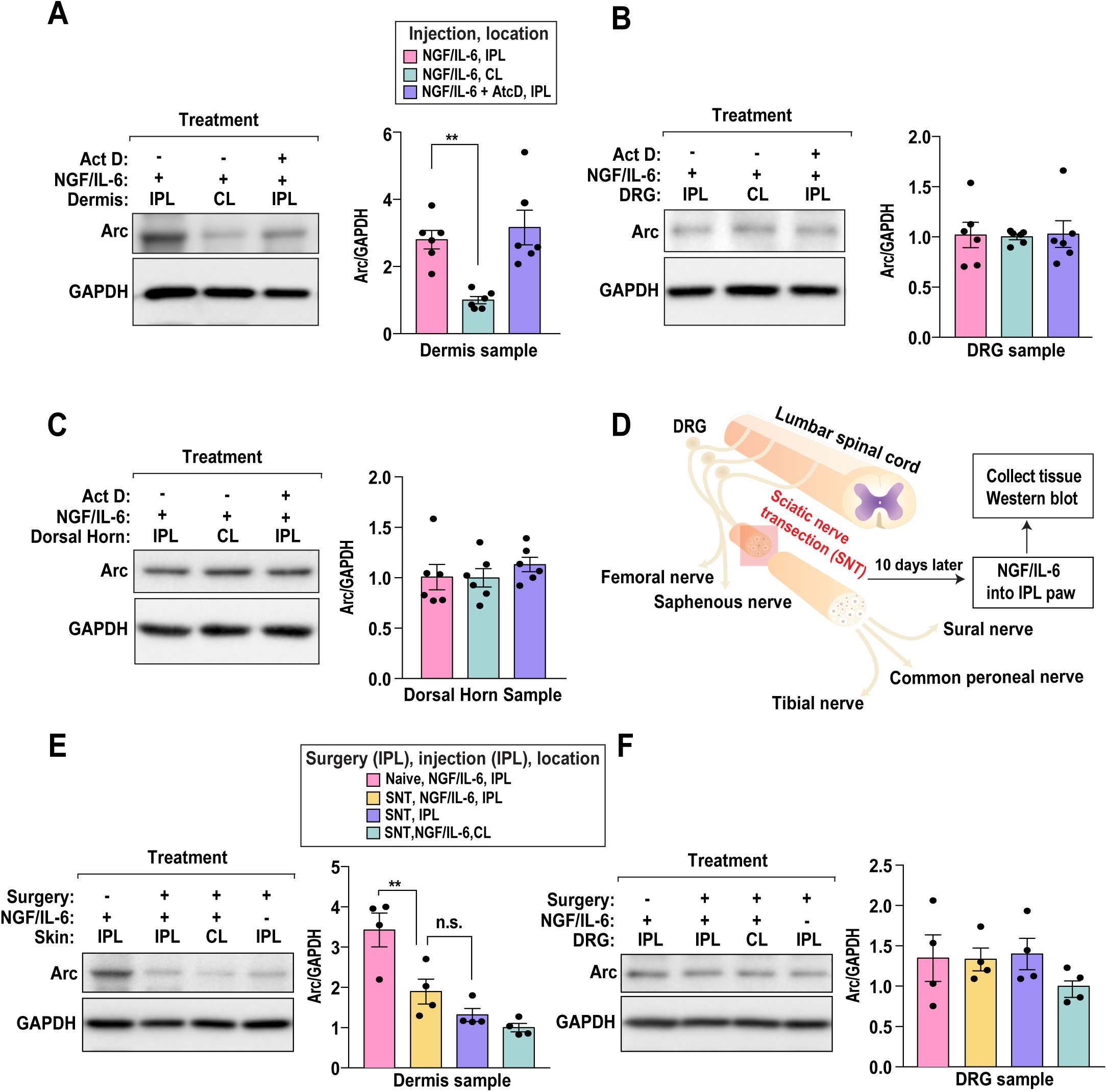
Induced biosynthesis of Arc in the skin depends on the presence of intact primary afferents. (A) Intraplantar administration of inflammatory mediators in the ipsilateral (IPL) paw of mice triggered a rapid translation of arc in skin that is not modify by the transcription inhibitor actinomycin D (Act D, 300 ng/25 μL). n = 6 animals per group. Ordinary one-way ANOVA: F_(2, 15)_ = 11.33, P = 0.0010. Bonferroni’s multiple comparisons test: NGF/IL-6, IPL vs NGF/IL- 6, CL; **P = 0.0022. IPL – ipsilateral or injected side, CL – contralateral or uninjected side. (B) Intraplantar administration of inflammatory mediators, in the presence of absence of Act D (300 ng/25 μL), did not alter Arc abundance in the DRG. (C) Intraplantar administration of inflammatory mediators, in the presence of absence of Act D (300 ng/25 μL), did not modify Arc abundance in the spinal dorsal horn (SDH). (D) A cartoon detailing the experimental protocol for sciatic nerve transection (SNT). Sciatic nerve was axotomized and ten days later, animals received an intraplantar injection of the inflammatory mediators. One hour post-injection, arc protein expression was analyzed in DRG and glabrous skin from injected (ipsilateral, IPL) and noninjected (contralateral, CL) paws. (E) Increase in Arc abundance in the skin caused by intraplantar administration of inflammatory mediators requires intact primary afferent fibers. n=4 animals per group. Ordinary one-way ANOVA: F_(3, 12)_ = 15.14, P = 0.0002. Tukey’s multiple comparisons test: Naïve, NGF/IL-6, IPL vs SNT, NGF/IL-6, IPL; **P = 0.0094. n.s. – non-significant. (F) SNT surgery transection did not alter Arc abundance in the DRG.

Arc is expressed in skin-migratory dendritic cells, thus, we sought to identify the cell type responsible for Arc production in the skin (Ufer et al., 2016). To determine if neurons are the relevant source, afferent fibers were eliminated through transection of the sciatic nerve on the ipsilateral side (Figure 3D). Transection of the sciatic nerve causes a complete loss of afferent fibers in the ipsilateral paw (Devor and Wall, 1981; Tandrup et al., 2000). After animals were allowed to heal for a period of ten days, NGF/IL-6 treatment was repeated. Arc expression was not increased as a result of the surgery. Injection of NGF/IL-6 on animals subjected to transection of the sciatic nerve no longer triggered an increase in Arc biosynthesis (Figure 3E). Additionally, transection of the sciatic nerve did not change Arc abundance in the DRG (Figure 3F). We conclude that NGF/IL-6 induced translation of Arc in the skin is likely due to local translation in afferent fibers.

### Arc regulates paw temperature in response to inflammation

Neurogenic inflammation plays a key role in vasodilation through release of neuropeptides from primary afferent fibers in response to noxious stimuli (Richardson and Vasko, 2002). Vasodilation increases blood flow to the skin which results in an increase in skin temperature (Chiu et al., 2012; Tanda, 2015). This increase in temperature can be detected by non-invasive forward looking infrared (FLIR) thermal imaging. To determine if Arc plays a role in neuroinflammation, we examined the temperature of the paw after a strong inflammatory challenge. We made use of a genetic model where a CreER reporter transgene induces polyadenylation prior to the Arc coding sequence (Guenthner et al., 2013). We found that homozygous animals have little remaining Arc in either the skin or the DRG (Figure S3). Because intraplantar injections of modest amounts of IL-6 and NGF do not elicit overt changes in the temperature of the paw, we made use of a more robust inflammatory mediator called Complete Freund’s Adjuvant (CFA) that increases levels of both NGF and IL-6 (Safieh-Garabedian et al., 1995; Su et al., 2012). Injection of CFA into the paw of mice results in a rapid increase in blood flow that manifested as an increase in temperature (Figure 4A). The average increase in paw temperature in wild-type animals was 1.4°C, across a 72-hour period. In contrast, lack of Arc resulted in a much higher increase in paw temperature, averaging at 2.6°C. This effect was significant at 3 and 24 hours after CFA administration (Figure 4B). Collectively, these data suggest that Arc is required to attenuate increases in skin temperature triggered by inflammation.

**Figure 4.**
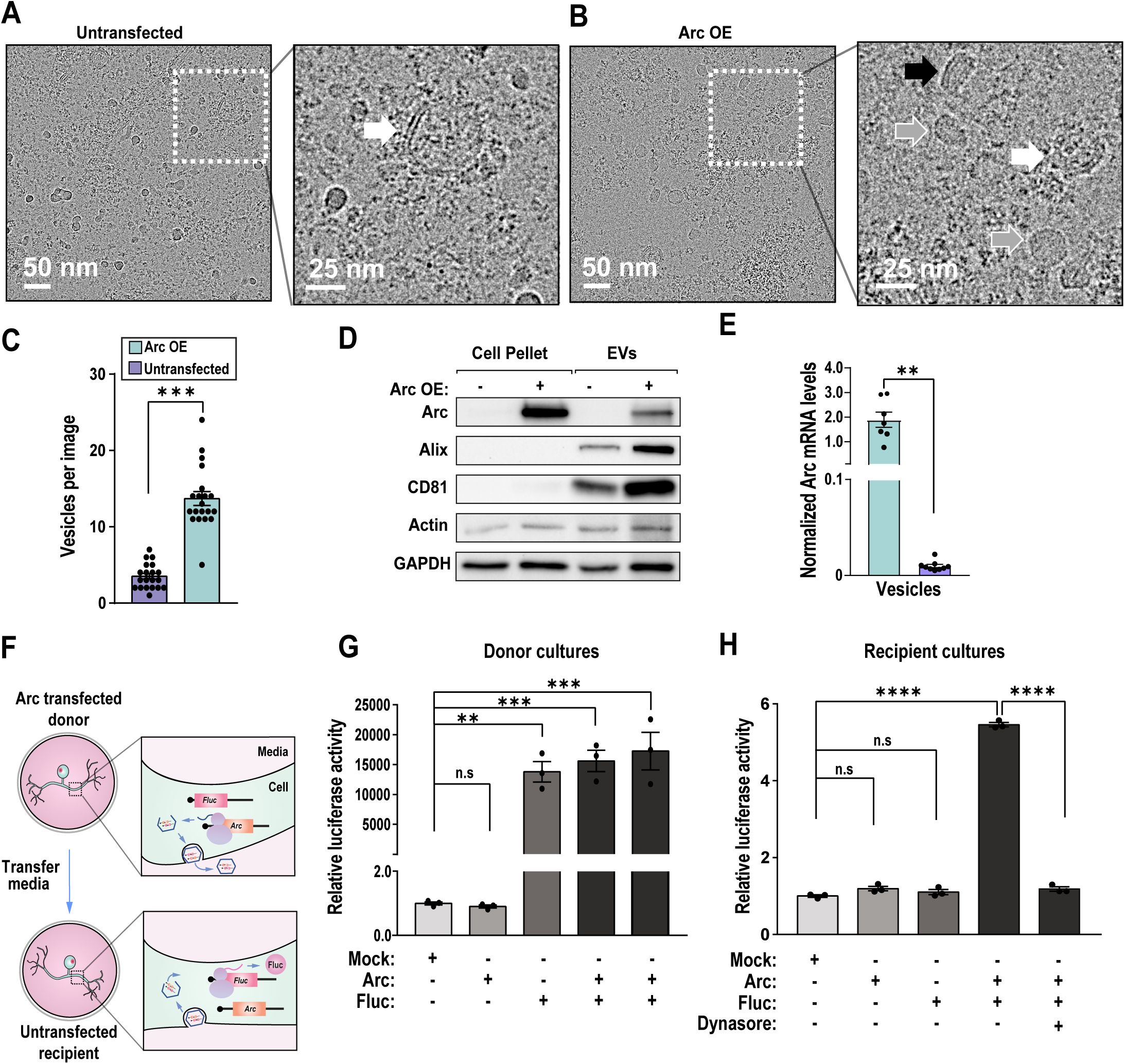
Sensory neurons produce *Arc*-containing extracellular vesicles. (A) Left panel, cryo-EM image of EVs purified from untransfected cells. Right panel, closeup of region with a membrane bilayer bound vesicle (white arrow). (B) Left panel, cryo-EM image of EVs purified from cells transfected with Arc. Right panel, closeup on three types of vesicles that increase in abundance. After transfection the most abundant vesicle type are single membrane vesicles (grey arrow). Low-density lipoprotein (LDL)-like vesicles (black arrow), which have a characteristic striped pattern due to stacked cholesteryl esters, and bilayered vesicles (white arrow) increase in abundance as well. (C) Left panel, average number of observed vesicles per image. 20 consecutive images for each sample were evaluated. The number of EVs/image increases ∼3.9 fold for the Arc transfected sample. Unpaired t test with Welch’s correction. ***P = 0.0003. (D) Immunoblot analysis of vesicles isolated from cells transfected with Arc are enriched for Arc, Actin, GAPDH and vesicular markers. (E) qPCR quantification of Arc mRNA reveals that Arc overexpression results in significant encashment. (n = 3). Unpaired t-test with Welch’s correction, **P = 0.0012. (F) A schematic depicting the model for Arc mediated RNA transfer. Media from cells that overexpress Arc may mediate transfer of abundant mRNA species such as firefly luciferase (*Fluc*). Transfer of media from donor cells to recipient cells may result in translation of encapsulated RNAs. (G) Firefly luciferase assay data for donor cultures normalized to a mock transfection sample. Ordinary one-way ANOVA: F_(4, 10)_ = 23, P = < 0.0001. Tukey’s multiple comparisons test: Mock vs Arc (n.s), P = > 0.999; Mock vs Fluc, **P = 0.0020; Mock vs Arc + Fluc (1), ***P = 0.0003; Mock vs Arc + Fluc(2), ***P = 0.0003. (H) Firefly luciferase assay data for recipient cultures normalized to a mock transfection sample. Ordinary one-way ANOVA: F_(4, 10)_ = 1220, P = < 0.0001. Tukey’s multiple comparisons test: Mock vs Arc (n.s), P = 0.1717; Mock vs Fluc (n.s), P = 0.6851; Mock vs Arc + Fluc, ****P = < 0.0001; Mock vs Arc + Fluc + Dynasore (n.s), P = 0.2012; Arc + Fluc vs Arc + Fluc + Dynasore, ****P = < 0.0001.

### Arc-containing extracellular vesicles can rescue inflammatory responses in Arc deficient mice

Given that Arc forms virus-like capsids that mediate mRNA transfer between different cell types, we hypothesized that Arc could facilitate communication between neurons and non-neuronal cells in the skin via EVs. Overexpression of Arc in HEK293 cell lines yields Arc-containing EVs capable of mRNA transport (Pastuzyn et al., 2018). To determine if nociceptors are capable of generating functional Arc vesicles, we made use of a nociceptor cell line derived from rat DRG neurons designated as F11 (Platika et al., 1985). Arc was expressed in F11 cells using a recombinant vector. EVs were enriched from bulk media using an EV isolation reagent (**STAR★Methods**). Afterward, dead cells were removed by low speed centrifugation. Vesicles were concentrated by ultracentrifugation and subjected to a gentle wash prior to resuspension. We subjected EV fractions to cryo-EM, then counted and characterized the morphology of the observed vesicles. Transfection of the F11 nociceptor cell line with Arc lead to an ∼3.9-fold increase of EVs per image (3.5 ± 1.7 vesicles/image in the control versus 13.7 ± 4.1 vesicles/image for transfected) (Figures 4C-E). EVs obtained from Arc-transfected cells were larger compared to the untransfected control (Figure S4). Thus, Arc expression in F11 cells induces the release of EVs that may contain Arc capsids.

As an additional test for the presence of Arc in the vesicles, we measured Arc protein release in EVs using Western blots (Figure 4F). Untransfected cells have little detectable endogenous Arc in either the cell pellet or in the EV fraction. In contrast, transfected cells show strong enrichment for Arc in the EV fraction. These EVs were highly enriched for known EV markers (e.g. ALIX, CD81, actin) (Andreu and Yanez-Mo, 2014; Willms et al., 2018). Arc-containing vesicles produced in HEK293 cells and cortical neurons show strong enrichment for the *Arc* mRNA (Pastuzyn et al., 2018). Using qPCR, we found that *Arc* mRNA can be robustly detected in EVs isolated from F11 cells expressing Arc (Figure 4G).

Arc vesicles mediate intercellular transfer of abundant mRNAs (Pastuzyn et al., 2018). As a functional test of Arc intercellular transfer obtained from EVs derived from nociceptors, we co-transfected Arc with a vector that encodes a firefly luciferase reporter (Figure 4H). Donor cells were transfected with Arc and the reporter, media was then replaced six hours later to remove residual complexes between DNA and transfection reagent. One day later, the media was collected and centrifuged to remove cell debris. The clarified media was placed on a recipient F11 culture and allowed to incubate for one day. Afterward, media was removed, and lysates were subjected to luminescence assays. We found that transfected cultures yielded substantial luciferase activity (Figure 4I). Transfer of the media from donor cells expressing luciferase reporter alone to non-transfected recipient cells resulted in little detectable activity (Figure 4J). In contrast, transfer of media from cultures co-expressing Arc and luciferase resulted in an increase in light production of 445 %. Transfer requires endocytosis in recipient cells as luciferase expression was diminished in cultures treated with a dynamin inhibitor (Dynasore). We conclude that Arc-containing EVs obtained from a nociceptor cell line are functionally similar to those obtained from HEK293 cells or primary hippocampal neurons (Pastuzyn et al., 2018).

To determine if Arc-containing EVs are able to rescue inflammatory responses in Arc deficient animals, we injected the ipsilateral paw with 1 µg of vesicles purified from either Arc overexpressing or non-transfected F11 cells and examined paw temperatures using forward looking infrared imaging or FLIR (Figures 5A). Injection of Arc-containing EVs rescued excessive heat responses in Arc KO animals at 3, 24, and 48 hours (Figure 5B). In contrast, injection of EVs obtained from non-transfected cells did not diminish the increase in hind paw temperature. Collectively, these data suggest that Arc EVs derived from nociceptor cells can restore normal functional responses in the skin of Arc KO mice

**Figure 5.**
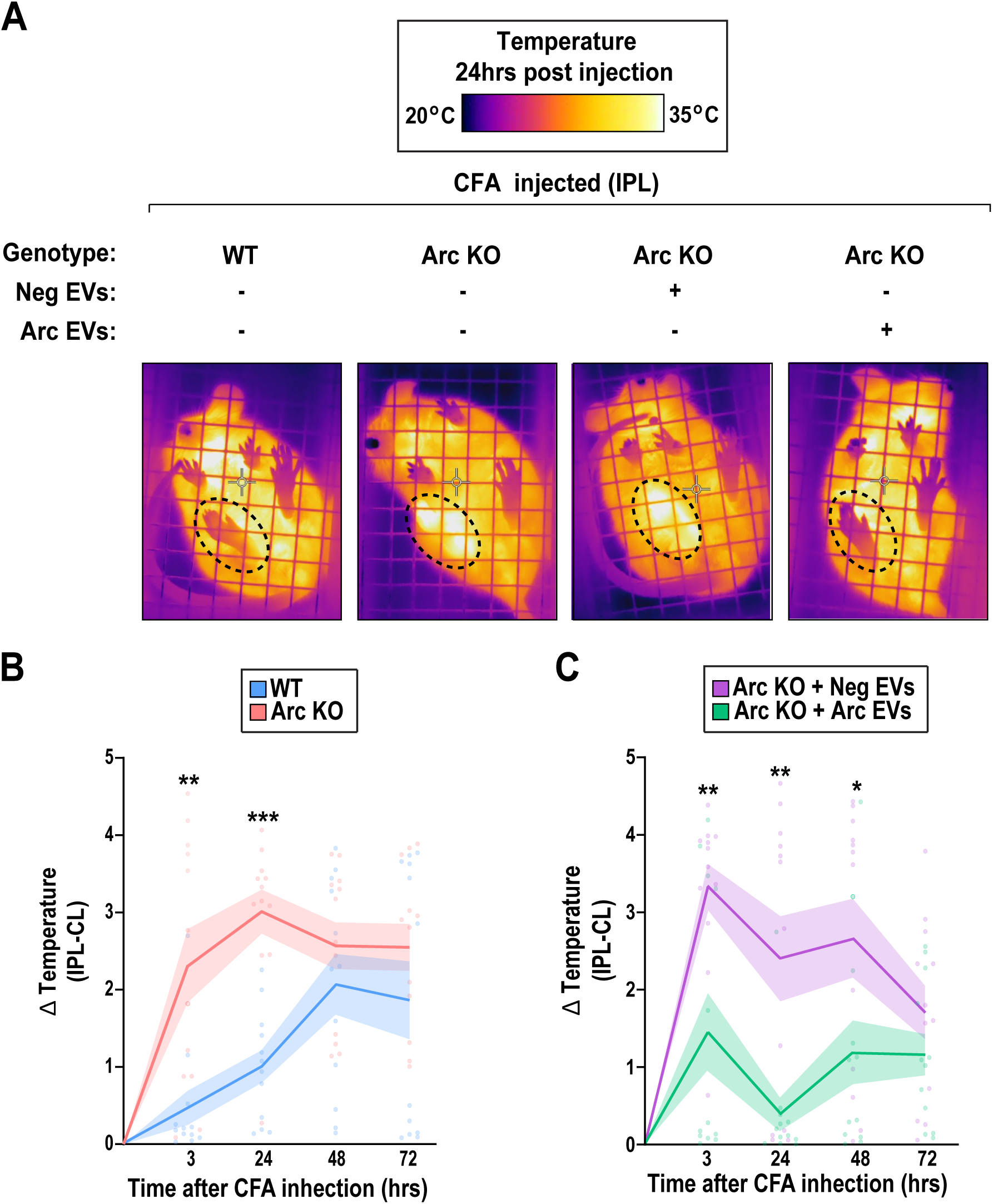
Arc-containing extracellular vesicles reduce thermal responses following injection of CFA. (A) Forward-looking infrared (FLIR) images of the paw of wild type (WT) and ARC knock-out (ARC KO) mice after injection of 5 µg of Complete Freund’s Adjuvant (CFA), with and without pretreatment of purified Arc vesicles (encircled paw). (B) Arc KO mice have increased ΔTemperature than WT mice, indicating exaggerated vasodilation. Two-way ANOVA: genotype effect, F(1, 88) = 25.62, P < 0.0001; time effect, F(3, 88) = 2.772, P = 0.0462. Bonferroni’s multiple comparison test: WT vs Arc KO, 3hrs **P = 0.0015, 24hrs ***P = 0.0005. Data are the difference in temperature between the injected (IPL) vs the uninjected (CL) paw of each mouse (ΔTemperature, °C). The thick line connects mean values and the flanking shaded areas denote S.E.M. n = 12 animals per group. (C) Pre-treatment with purified Arc containing EVs rescues the exaggerated vasodilation in the Arc KO mice. Two-way ANOVA: vesicle effect, F_(1, 88)_ = 26.75, P < 0.0001; time effect, F_(3, 88)_ = 2.720, P = 0.0493. Bonferroni’s multiple comparison test: WT vs Arc KO, 3hrs **P = 0.0057, 24hrs **P = 0.0027, 48hrs **P = 0.0458. As in panel B, the data represent change in temperature between IPL and CL paws. The thick line connects mean values and the flanking shaded areas denote S.E.M. n = 12 animals per group

We wondered if the cause of the increased thermal responses to CFA was driven by vasodilation. As a measure of blood flow, we made use of 3-dimensional (3D) high-frequency ultrasound imaging. In these assays blood flow is measured using a power Doppler method. We found that the relative blood flow in paws was higher in Arc animals relative to wild-type mice (Figure 6A). However, the amount of swelling was not significantly different between groups at 24 hours (Figure 6B, left). To determine if Arc EVs attenuate the increase in blood flow, we injected Arc deficient animals (Figure 6B, right). We found that the Arc-containing EVs recued abnormal vasodilation whereas the negative control EVs did not normalize elevated blood flow. When taken together with the FLIR measurements, our data suggest that intercellular Arc signaling constrains neurogenic inflammation in the periphery.

**Figure 6.**
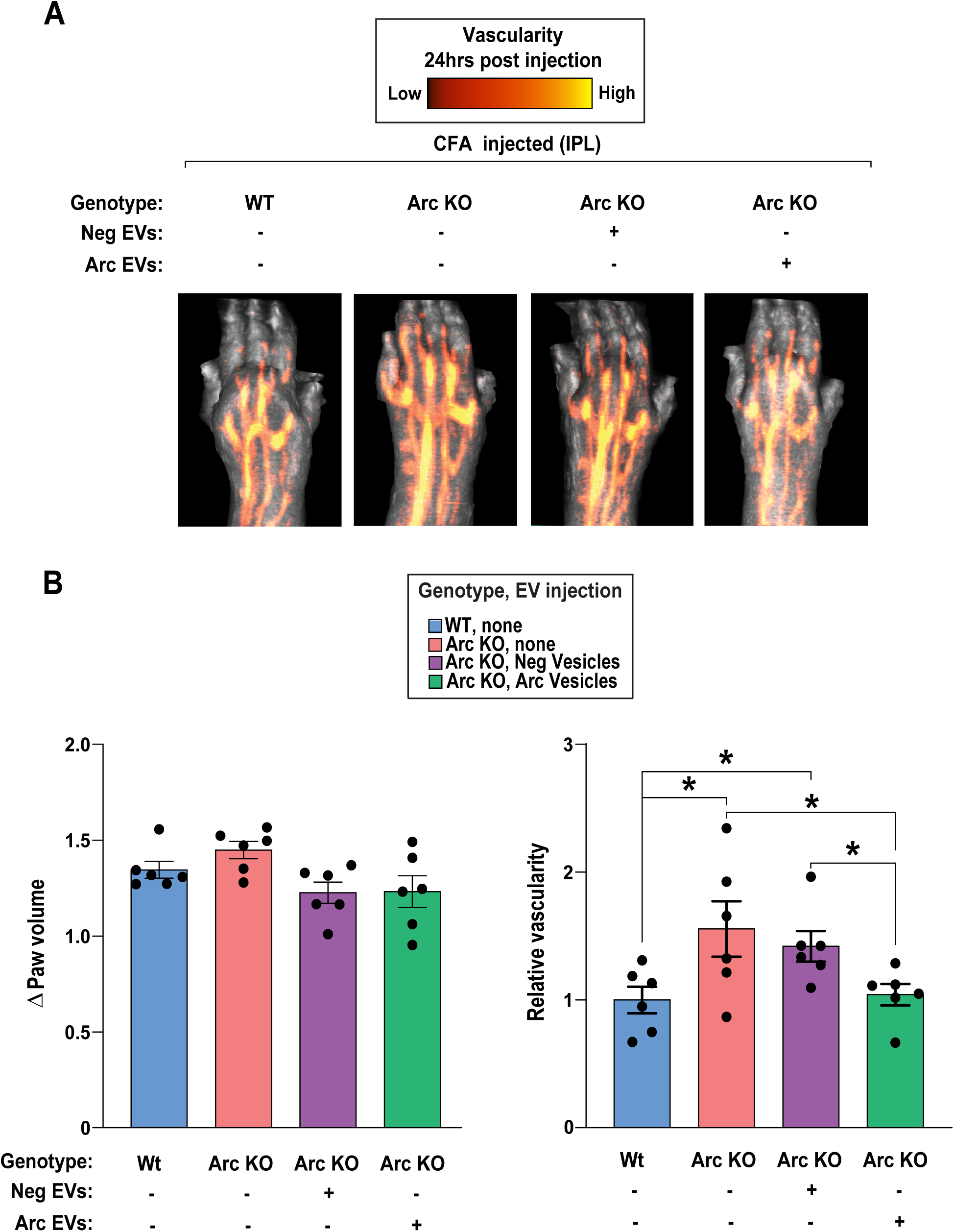
Arc-containing extracellular vesicles reduce vascularity in the skin following injection of CFA. (A) Doppler ultrasound images of the paw of wild type (WT) and ARC knock-out (ARC KO) mice after injection of 5 µg of Complete Freund’s Adjuvant (CFA). Vesicle treatments were either from cells expressing Arc or an vesicles isolated from an transfected negative control. (B) There was no difference in paw volumes of WT and Arc KO mice, with and without pretreatment of purified Arc vesicles, after CFA injection. Each bar represents the mean ± S.E.M. of the change in volume between the CFA-injected and uninjected paw (ΔPaw volume). n = 6 animals per group. (C) Arc KO mice have increased vascularity than WT mice after CFA-injection, indicating exaggerated vasodilation. Pre-treatment with purified Arc vesicles, but not of Neg vesicles, rescues the exaggerated vasodilation in the Arc KO mice. Unpaired t-test, WT vs Arc KO *P = 0.0432, WT vs Arc KO + Neg EV *P = 0.0244, ARC KO vs ARC KO + Arc EV *P = 0.0258, Arc KO + Neg EV vs Arc KO + Arc EV *P = 0.0276. Each bar represents the mean ± S.E.M. of the relative change in percent vascularity after CFA injection. n = 6 animals per group.

## DISCUSSION

We describe a novel role for intercellular Arc signaling in nociceptor mediated neuroinflammatory responses in the skin. The unexpected finding that Arc mediates cell-to-cell communication similar to retrovirus “infection” has prompted speculation as to what kind of functional role this pathway may control (Kedrov et al., 2019). We posit that translation of Arc in DRGs in response to inflammatory signals, leads to the generation and release of Arc EVs that regulate inflammation in the skin. Furthermore, our observations have broad implications for the identification of targets of translational control in sensory neurons. The ability to precisely control treatments consisting of inflammatory mediators coupled to global measurement of nascent protein synthesis affords unique opportunities to identify targets of preferential translation. This work complements related efforts to profile translation in tissues (Megat et al., 2019; Uttam et al., 2018). We focused on identification of transcripts that display rapid changes in translation as opposed to regulatory changes that manifest days after an injury. Our reasoning was that widespread changes in transcription were likely to be minimal after only 20 minutes. Indeed, the number of transcripts with significant changes in translation is nearly double that of those whose levels change in abundance (Figure S1B). We identified multiple immediate early genes (*e*.*g*. Egr2, Fos) that were preferentially translated in response to a brief treatment with inflammatory mediators. We focused specifically on Arc because it is rapidly transcribed in the brain and localized to dendrites prior to local translation and because of its recently established roles related to intercellular signaling (Ashley et al., 2018; Link et al., 1995; Lyford et al., 1995; Pastuzyn et al., 2018; Steward et al., 1998; Wallace et al., 1998). Disruption of Arc through knockdown or genetic ablation results in profound deficits in learning and memory (Guzowski et al., 2000; Plath et al., 2006). Yet, Arc remains poorly characterized in sensory neurons. Our observations enable three major conclusions.

Our experiments uncover a new function for Arc in the periphery. We found that Arc plays a role in inflammation. Two of the best characterized nociceptive transmitters are CGRP and substance P (Brain and Williams, 1989; Saria, 1984). Both are potent vasodilators released from sensory fibers in response to noxious cues. Intriguingly, Arc appears to function in an opposing manner. Why would different subsets of sensory neurons produce signaling molecules with opposing effects on the same process? Transcriptional control of the lipopolysaccharide (LPS) response in macrophage provides a relevant parallel. LPS induces rapid transcription of genes that encode functionalities linked to cell migration and tissue repair (Medzhitov and Horng, 2009). Intriguingly, LPS also stimulates production of inducible negative regulators, such as IκBα, that limit inflammation by interfering with pro-inflammatory transcription factors (Kuwata et al., 2006; Wessells et al., 2004). Negative feedback loops are broadly critical in immunity as they mitigate deleterious pathophysiological consequences that arise from excessive inflammation. We propose that this axiom also extends to inflammatory signaling driven by sensory neurons.

Second, we establish that Arc is translated in afferent fibers in the skin. The dermis contains a diverse microenvironment that includes nerve fibers, keratinocytes, fibroblasts, nerve terminals, basal cells, capillaries, and resident immune cells. Non-neuronal cells collaborate to control the function of sensory neurons through multiple mechanisms. For example, keratinocytes facilitate detection of mechanical cues through release of ATP (Moehring et al., 2018). Immune cells secrete a range of molecules that sensitize nociceptive neurons (*e*.*g*. NGF, IL-6, PGE2 etc.) (Pinho-Ribeiro et al., 2017). Communication from sensory neurons to other cell types is mediated almost exclusively by peptides. We found that Arc deficient mice display exaggerated vasodilatation that can be rescued by injection of purified vesicles containing Arc. Based on these observations, we propose a model where potential intercellular transport of RNA mediates communication between sensory neurons and epithelial cells. We show that purified Arc-containing vesicles obtained from a nociceptor cell line are enriched for *Arc* mRNA. However, the vesicles likely contain multiple RNA species. While our data do not discount the possibility that the Arc protein itself could be the relevant molecule responsible for rescue, it is clearly of interest to determine the RNA content of the vesicles with the overarching goal of understanding the anti-inflammatory effects of the purified vesicles.

Third, Arc is likely translated from existing mRNA. In the brain, synaptic activity results in transcription and localization of Arc to sites of local protein synthesis (Steward et al., 1998). We found that introduction of transcriptional inhibitors did not prevent Arc accumulation in the dermis. This suggests that Arc is primarily translated from existing mRNA which departs from compelling data obtained in the rat hippocampus (Farris et al., 2014). The use of slightly different mechanisms may reflect a key anatomical challenge to local translation – distance. Sensory neurons possess the longest axons in the body that can extend for a meter in humans. The rate an mRNA travels is at best approximately 5 μm/s (Park et al., 2014). Thus, a reasonable estimate for the amount of time required to transport a newly transcribed RNA a meter is 55 hours. Local translation is an attractive mechanism to achieve Arc biosynthesis on demand. It is unclear how Arc is maintained in a quiescent state in fibers. One potential mechanism is through post-translational modifications of translation factors that favor preferential patterns of translation. For example, mGluR activation stimulates rapid translation of Arc through Eukaryotic Elongation Factor 2 Kinase (eEF2K) dependent phosphorylation of Eukaryotic Elongation Factor 2 (eEF2) (Park et al., 2008). This in turn reduces elongation rates of most transcripts. By an unknown mechanism, Arc evades repression by phosphorylated eEF2 in dendrites. It would be of interest to determine if similar regulation occurs in afferent fibers.

To summarize, translational control is prominent in sensory neurons (de la Pena et al., 2019; Loerch et al., 2019). Beyond the established role of *de novo* protein synthesis in pain associated behaviors, our experiments uncover a new target of translational regulation – Arc. We find that Arc is expressed throughout the DRG and accumulates in the skin in response to inflammatory mediators. The accumulation requires afferent fibers and is insensitive to transcriptional inhibition. We find that genetic disruption of Arc results in exaggerated inflammation that is normalized upon introduction of extracellular vesicles containing Arc. This suggests that Arc is a new type of anti-inflammatory mediator. Given parallel negative feedback in other systems, this is important for understanding the process of neuroinflammation. Indeed, activity dependent vasodilation also occurs in the brain (Chow et al., 2020). Our data suggest that virus-like intercellular communication is a key regulator of neuron to non-neuronal signaling in the periphery and reveal a novel mechanism for Arc-dependent regulation of neuroinflammation.

## STAR★METHODS

### Contact for reagents and resource sharing

Further information and requests for resources and reagents should be directed to and will be fulfilled by the Lead Contact Zachary Campbell.

## Key Resources Table

| REAGENT or RESOURCE | SOURCE | IDENTIFIER |
| --- | --- | --- |
| **Antibodies** |  |  |
| Rabbit monoclonal anti-GAPDH(14C10) | Cell Signaling Technology | Cat#2118 |
| Mouse monoclonal anti-Arc (C-7) | Santa Cruz Biotechnology | Cat#sc-17839 |
| Goat anti Mouse IgG, Alexa Fluor 405 | Thermo Fisher Scientific | Cat#A31553 |
| Mouse monoclonal anti-Neurofilament 200 kDa (N52) | MilliporeSigma | Cat#MAB5266 |
| Goat anti-mouse IgG, light chain specific | Jackson ImmunoResearch Laboratories | Cat#115-035-174 |
| Goat Anti-Rabbit IgG (H+L) | Jackson ImmunoResearch Laboratories | Cat#111-036-144 |
| Mouse monoclonal anti-Alix (3A9) | Cell Signaling Technology | Cat#2171 |
| Rabbit monoclonal anti-CD81 (D5O2Q) | Cell Signaling Technology | Cat#10037 |
| Rabbit polyclonal anti-β Actin | Cell Signaling Technology | Cat#4967 |
| Rabbit monoclonal anti-GFP | Cell Signaling Technology | Cat#2956 |
| Chicken polyclonal anti-peripherin | Novus Biologicals | Cat#NBP1-05423 |
| **Chemicals, Peptides, and Recombinant Proteins** |  |  |
| DMEM/F-12, GlutaMAX™ Supplement | Gibco | Cat#10565018 |
| Hank's Balanced Salt Solution | Gibco | Cat#14170112 |
| Collagenase A | Roche | Cat#10103578001 |
| Collagenase D | Roche | Cat#11088858001 |
| Papain | Roche | Cat#10108014001 |
| Trypsin Inhibitor | Roche | Cat#10109886001 |
| Bovine Serum Albumin | Thermo Fisher Scientific | Cat# BP1600-100 |
| Fetal Bovine Serum - Premium Select | R&D Systems | Cat#S11550 |
| 5-Fluoro-2'-deoxyuridine | Sigma-Aldrich | Cat#F0503 |
| Uridine | Sigma-Aldrich | Cat#U3003 |
| Poly-D-lysine hydrobromide | Sigma-Aldrich | Cat#P7886 |
| Mouse Nerve Growth Factor 2.5S Protein | MilliporeSigma | Cat#01-125 |
| Recombinant Mouse IL-6 Protein | R&D Systems | Cat#406-ML-005 |
| Emetine, Dihydrochloride | MilliporeSigma | Cat#324693 |
| RNase-Free DNase I | Lucigen Corporation | Cat#D9905K |
| RNasin Ribonuclease Inhibitors | Promega Corporation | Cat#N2111 |
| Prostaglandin E2 | Cayman Chemical | Cat#14010 |
| Normal Goat Serum | Sigma-Aldrich | Cat# NS02L |
| DAPI (4',6-Diamidino-2-Phenylindole, Dihydrochloride) | Thermo Fisher Scientific | Cat#D1306 |
| ProLong Gold Antifade Mountant | Thermo Fisher Scientific | Cat#P36930 |
| Protease Inhibitor Cocktail powder | Sigma-Aldrich | Cat#P2714 |
| Phosphatase Inhibitors Cocktail 2 and Cocktail 3 | Sigma-Aldrich | Cat#P5726 and P0044 |
| Total Exosome Isolation Reagent | Thermo Fisher Scientific | Cat# 4478359 |
| Lipofectamine 3000 Transfection Reagent | Thermo Fisher Scientific | Cat# L3000008 |
| Promega ImProm-II™ Reverse Transcription | Thermo Fisher Scientific | Cat# PRA3800 |
| iQ SYBR Green Supermix | Bio-Rad Laboratories | Cat# 1708886 |
| **Critical Commercial Assays** |  |  |
| Western Chemiluminescent HRP Substrate (ECL) | MilliporeSigma | Cat# WBKLS0500 |
| SMARTer smRNA-Seq kit for Illumina | Takara Bio USA, Inc | Cat#635031 |
| QuantSeq 3' mRNA-Seq Library Prep Kit FWD for Illumina | Lexogen | Cat#015.96 |
| RiboCop rRNA Depletion Kit V1.2 | Lexogen | Cat#037.96 |
| Qubit dsDNA HS Assay Kit | Thermo Fisher Scientific | Cat#Q32851 |
| High Sensitivity NGS Fragment Analysis Kit | Advanced Analytical Technologies, Inc | Cat#DNF-474 |
| NextSeq 500/550 High Output v2 kit (75 cycles) | Illumina | Cat# FC-404-2005 |
| RNAscope Protease III & Protease IV Reagents kit | Advanced Cell Diagnostics | Cat#322340 |
| RNAscope Multiplex Fluorescent Detection Kit | Advanced Cell Diagnostics | Cat#320851 |
| RNAscope Wash Buffer Reagents | Advanced Cell Diagnostics | Cat#310091 |
| RNAscope Probe - Mm-Arc | Advanced Cell Diagnostics | Cat#316911 |
| RNAscope Probe - Mm-Calca-alltv-C2 | Advanced Cell Diagnostics | Cat#417961-C2 |
| RNAscope Probe - Mm-P2rx3-C3 | Advanced Cell Diagnostics | Cat#521611-C3 |
| **Deposited Data** |  |  |
| Sequencing deposited to GEO | https://www.ncbi.nlm.nih.gov/geo/query/acc.cgi?acc=GSE117043 | GSE117043<br>(Token for access: efgpqousplafvif ) |
| **Experimental Models: Organisms/Strains** |  |  |
| Swiss Webster Mice | Taconic laboratories | Model# SW-M |
| C57BL/6J Mice | Taconic laboratories | Model# B6-M |
| B6.129(Cg)-Arc<sup>tm1.1(cre/ERT2)Luo/J</sup> Mice | Jackson Laboratory | Cat# 021881 |
| **Oligonucleotides** |  |  |
| Primers | Sigma |  |
| **Recombinant DNA** |  |  |
| PLJM1 - Arc | This manuscript |  |
| PGL4.13[luc2/SV40] | Promega | Cat# E6681 |
| **Software and Algorithms** |  |  |
| TopHat 2.1.1 | https://ccb.jhu.edu/software/tophat/index.shtml |  |
| Bowtie 2.2.9 | http://bowtie-bio.sourceforge.net/bowtie2/index.shtml |  |
| Cufflinks 2.2.1 | http://cole-trapnell-lab.github.io/cufflinks/ |  |
| SerialEM | https://bio3d.colorado.edu/SerialEM/ | (Mastronarde, 2005) |
| cisTEM | https://cistem.org/ | (Grant et al., 2018) |
| Tdisp | https://sourceforge.net/projects/tigris/ |  |

### Experimental Model and Subject Details

*In vitro* experiments were performed using wild-type male Swiss Webster and C57BL/6J mice (4 weeks old) purchased from Taconic laboratories. Behavioral experiments were performed using male Swiss Webster, C57BL/6J and Arc^CreER^ mice (Jackson stock #021881) between the ages of 8 and 12 weeks, weighing approximately 25 – 30 g at the start of the experiment. Animals were housed with a 12 hr light/dark cycle and had food and water available *ad libitum*. All animal procedures were approved by Institutional Animal Care and Use Committee at The University of Texas at Dallas and were in accordance with International Association for the Study of Pain guidelines.

## METHOD DETAILS

### Ribosome profiling cultures

Dorsal root ganglia (DRG) from cervical (C1) to lumbar (L6) spinal segment were excised from 10 mice per replicate and placed in chilled Hanks’ balanced salt solution (HBSS; Invitrogen). Following dissection, DRGs were enzymatically dissociated with collagenase A (1 mg/mL, Roche) for 25 min and collagenase D (1 mg/mL, Roche) with papain (30 U/mL, Roche) for 20 min at 37 °C. DRGs were then triturated in a 1:1 mixture of 1 mg/mL trypsin inhibitor (Roche) and bovine serum albumin (BioPharm Laboratories), then filtered through a 70 μm cell strainer (Corning). Cells were pelleted, then resuspended in DRG culture media: DMEM/F12 with GlutaMAX (Thermo Fisher Scientific) containing 10% Fetal Bovine Serum (FBS; Thermo Fisher Scientific), 1% penicillin and streptomycin, and 3 μg/mL 5-fluorouridine with 7 μg/mL uridine to inhibit mitosis of non-neuronal cells. Cells were evenly distributed in three poly-D-lysine-coated culture dishes (100 mm diameter) (BD Falcon) and incubated at 37 °C in a humidified 95% air/5% CO_2_ incubator. DRG culture media was changed every other day and cells were treated with NGF (20 ng/mL) and IL-6 (50 ng/mL) at day 6 for 20 minutes followed by addition of emetine (50 μg/mL) for 1 minute in order to protect the ribosome footprints.

### Library generation and sequencing

Libraries consisting of ribosome bound RNA fragments were generated as described with minor adjustments in the composition of the polysome lysis buffer (20 mM Tris-HCl, pH 7.5, 250 mM NaCl, 15 mM MgCl_2_, 1 mM DTT, 0.5% (vol/vol) Triton X-100, 2.5 U/ml DNase I, 40 U/mL RNasin and 50μg/mL emetine) (Hornstein et al., 2016; Ingolia, 2010). MicroSpin S-400 columns (GE Healthcare) were used to isolate ribosome bound RNAs. After rRNA was removed using RiboCop rRNA depletion kit (Lexogen), footprints were dephosphorylated then size selected (28-34nt) by PAGE on 15% TBE-Urea gels (Bio-Rad). Footprints were generated by SMARTer smRNA-Seq kit for Illumina (TaKaRa). RNA abundance was quantified using the Quantseq 3′ mRNA-Seq library kit (Lexogen). The concentrations of purified libraries were quantified using Qubit (Invitrogen) and the average size was determined by fragment analyzer with high sensitivity NGS fragment analysis kit (Advanced Analytical Technologies Inc.). Libraries were then sequenced on an Illumina NextSeq500 sequencer using 75-bp single-end high output reagents (Illumina).

After sequencing, files were downloaded from a BaseSpace onsite server. An Initial quality check was conducted using FastQC 0.11.5 (Babraham Bioinformatics). Adapters were subject to trimming based on adapter sequences. Mapping was conducted with TopHat 2.1.1 (with Bowtie 2.2.9) to the mouse reference genome (NCBI reference assembly GRCm38.p4) and reference transcriptome (Gencode vM10). Strand orientation was considered during the mapping process. Processed bam files were quantified for each gene using Cufflinks 2.2.1 with gencode.vM10 genome annotation. Read counts were not normalized by length by the using the Cufflinks option -- no–length–correction. Relative abundance for the i^th^ gene was determined by calculating TPM (transcripts per million) values as follows:

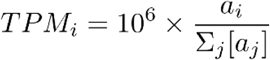, where a_j_ is the Cufflinks reported relative abundance.

Finally, TPM values were normalized to upper decile for each biological replicate and udTPM (upper decile TPM) were used for analysis (Glusman et al., 2013),to provide uniform processing for samples with varying sequencing depth.

### Single cell data

Single cell DRG sequencing data were generated based on published data (Li et al., 2016). Seurat package 2.2.1 (Butler et al., 2018) was used for visualization (Van der Maaten and Hinton, 2008).

### Immunofluorescence, immunohistochemistry, and *in situ* hybridization

Spinal cord and L4-L5 DRG tissues were removed from adult 8- to 12-week-old mice and mounted using OCT mounting medium. Tissues were cryosectioned to 20 μm thickness and mounted onto Superfrost slides (Thermo Fisher Scientific). The sections were subsequently immersed in prechilled 10% formalin for 15 min and dehydrated in increasing concentration of ethanol (50%, 70% and 100%). After complete drying, each section was carefully circled using a hydrophobic barrier pen (Immedge pen, RNAscope, ACDBio) and treated with Protease IV (RNAscope Protease III & Protease IV Reagents kit, ACDBio) at 40°C for 2 min. The sections were then washed twice in PBS and target probes that can be detected in three different color channels respectively (Atto 550, Alexa 488 and Atto 647) were added according to the manufacturer’s protocol (RNAscope Fluorescent Multiplex Reagent kit). Briefly, the Arc (Arc, accession no. NM_018790.2, 20 bp, target region: 23-1066), CGRP (CGRP, accession no. NM_007587.2, 17 bp, target region: 44-995) and P2×3 probes (P2×3, accession no. NM_145526.2, 20 bp, target region: 795-1701) were added and incubated at 40°C for 2 hr. Afterward, the sections were washed in RNAscope wash buffer twice and treated with the amplification reagent 1 (AMP1) for 30 min. There were a total of 4 amplification reagents (AMP1-AMP4) with alternating washes of 30 and 15 min incubation period. After the RNAscope protocol was completed, slides were blocked for 1 hr in 10% normal goat serum (NGS), 0.3% Triton X-100 in 0.1M PB. DRG slices were incubated overnight with a primary antibody against NF200 (1:500, Millipore, cat # MAB5266). The next day, slides were incubated with Goat anti-Mouse Alexa Fluor 405 (1:1000, ThermoFisher Scientific, A-31553) for 1 hr. After additional PBS washes, slides were coverslipped with Prolong Gold antifade and allowed to cure for 24 hr before imaging.

For immunohistochemistry, DRG and sciatic nerve sections were fixed in ice-cold 4% paraformaldehyde in 1× TBS for 1 h and then subsequently washed three times for 5 min each in 1× TBS. Slides were then incubated in a permeabilization solution made of 1× TBS with 0.4% Triton X-100 (Sigma-Aldrich) for 30 min and then were washed three times with 1× TBS. Tissues were blocked for at least 2 h in 10% heat-inactivated NGS in 1× TBS. Primary antibodies were used to detect the following proteins: GFP (1:200, CST, cat. # 2956,), peripherin (1:1000; Novus, cat. # NBP1-05423, NF200 (1:500, Millipore, cat # MAB5266). Primary antibodies were applied and incubated at 4 °C overnight. The next day, slides were washed with 1x TBS and then appropriate secondary antibodies (Alexa Fluor, Invitrogen) were applied for 2 h. After additional washes, coverslips were mounted on slides with ProLong Gold antifade mountant (Invitrogen).

All of the images corresponding to immunohistochemistry and *in situ* hybridization samples were acquired using an FV-3000 confocal microscope (Olympus). For each DRG image, colocalization of *Arc*/*Calca* (CGRP) and Arc/*P2×3* (P2×3) was calculated using ImageJ plug-in JACoP (Bolte and Cordelieres, 2006) and represented as % of Arc-positive cells expressing markers (*Calca or P2×3*). Colocalization (*Arc*/NF200) was assessed by counting the number of *Arc*-positive neurons that showed obvious *Arc* mRNA expression within the soma.

### Protein immunoblotting

For *in vitro* experiments, Swiss Webster mice (4-week-old) were anesthetized with isoflurane and euthanized by decapitation. The DRGs from C1 to L6 spinal column were dissected and placed in chilled HBSS solution. Immediately after dissection, DRGs were dissociated and triturated as described above in the ribosome profiling culture. After dissociation, cells from five mice were distributed evenly in a 6-well plate coated with poly-D lysine and maintained at 37°C in a humidified 95% air/5% CO_2_ incubator with fresh culture media replacement every other day. At day 6, the cells were treated with NGF (20 ng/mL) and IL-6 (50 ng/mL) for 20, 60, and 120 minutes. Cells were rinsed with chilled PBS then lysed in lysis buffer (50 mM Tris, pH 7.4, 150 mM NaCl, 1 mM EDTA, pH 8.0, and 1% Triton X-100) containing protease and phosphatase inhibitors (Sigma-Aldrich). Total protein was extracted by ultrasonication and the supernatant was collected by centrifugation at 18,000 × g for 15 min at 4 °C. Protein samples in 1× Laemmli sample buffer (Bio-Rad) were loaded and separated by 10% SDS-PAGE gels and then transferred to Immobilon-P membranes (Millipore). Membranes were blocked in Tris Buffered Saline (TBS) with 5% low-fat milk for 2 hr at room temperature, then incubated overnight at 4°C with primary antibodies against Arc (1:300, Santa Cruz Biotechnology, cat # sc-17839), Alix (1:1000, Cell Signaling Technology, cat # 2171) or CD81(1:1000, Cell Signaling Technology, cat # 10037). After primary incubation, membranes were washed in TBST (TBS with 0.05% Tween 20) for 30 min (3 × 10 min) and incubated with secondary antibodies conjugated to horseradish peroxidase (Jackson ImmunoResearch) for 1 hr at room temperature. The signal was detected using Western Chemiluminescent HRP Substrate (ECL) (MilliporeSigma) on ChemiDoc Touch Imaging System (Bio-Rad). The blot was stripped in Restore Plus western blot stripping buffer (Thermo Fisher) for 25 min and reincubated with primary antibody against GAPDH (1:10,000, Cell Signaling Technology, cat # 2118) or Actin (1:10,000, Cell Signaling Technology, cat # 4967) for 1 hr at room temperature as an internal control. Membranes were washed 3 × 10 min in TBST and incubated with a secondary antibody for GAPDH or Actin. For the *in vivo* experiments, ipsilateral and contralateral paws, spinal cords and L4-L5 DRGs from adult 8- to 12-week-old C57BL/6J mice were rapidly removed and lysed in cell lysis buffer. Total protein was collected as described above. Analysis was performed using Image Lab 6.0.1 (Bio-Rad). Phosphorylated proteins were normalized to their respective total proteins and expressed as a percent of change compared to vehicle groups.

### Sciatic nerve transection

For transection experiments, the left sciatic nerve of 8- to 12-week-old C57BL/6J mice was exposed and transected at midthigh level under isoflurane anesthesia. The wound was closed with 5-0 silk sutures. Following surgery, animals were housed and allowed to recover for 10 days before intraplantar NGF/IL-6 in the presence or absence of actinomycin D (Act D) administration.

### Cloning

The PLJM1 – eGFP vector (Addgene) was digested with NheI (Thermo Fischer Scientific) and EcoRI (Thermo Fischer Scientific) to excise the eGFP sequence. Mouse Arc CDS (Ensembl: ENSMUST00000023268.13) was cloned using the primers (5’ ATAAGCAGAGCTGGTTTAGTGAACCGTCAGATCCGCTAGCGCCACCATGGAGCTGGACCATATGACC3’ and 5’ TGTGGATGAATACTGCCATTTGTCTCGAGGTCGAGAATTCCTATTCAGGCTGGGTCCT G 3’ from mouse brain cDNA into PLJM1 using Gibson assembly (Gibson et al., 2009).

### F11 Transfection

F11 cells were grown in DMEM/F-12 media (Thermo Fisher Scientific) supplemented with 10% FBS and 1% Penicillin and Streptomycin at 37°C at 5% CO2. Cells were transfected with the Arc overexpression vector at a confluency of 70% using lipofectamine 3000 (Thermo Fisher Scientific). Transfection media was replaced with DMEM/F12 supplemented with 10% exosome free FBS and 1% Penicillin and Streptomycin after 6 hr. Cells were washed twice with 1X PBS before media replacement. After 24 hr, media from transfected cell lines was harvested for extracellular vesicle Isolation and downstream assays.

### Extracellular vesicle isolation

Extracellular vesicles were isolated using Total Exosome Isolation Reagent (Thermo Fisher Scientific). Harvest media was spun down at 2000 x g for 30 mins to remove cell debris. The supernatant was mixed with the isolation reagent in a ratio of (2:1) and incubated at 4°C for 16 hr. After incubation, samples were centrifuged at 10,000 x g for 1 hr to obtain a crude EV pellet. The pellet was resuspended in cold 1X PBS and centrifuged at 100,000 x g for 70 minutes to wash out contaminants. The final pellet was resuspended in cold 1 X PBS and used for downstream applications.

### Vesicle Exchange Assay

24 hours following transfection, media was aspirated from donor cultures and centrifuged 2000 x g for 10 minutes. Media was aspirated again and centrifuged at 3000 x g for 20 minutes to remove cell debris. Media was removed from recipient cells and the donor culture supernatant was transferred onto recipient cells and incubated for 24 hours at 37°C and 5%CO_2_. Cultures treated with dynamin inhibitors were incubated with 80µM Dynasore concurrently with media transfer. After 24 hours, the cells were washed with 1X PBS and then lysed with 1X Passive Lysis Buffer (Promega) for 10 mins at 4°C. Firefly luciferase activity was assessed using LARII substrate (Promega). To 100µl of cell lysate was mixed with 50 µl of substrate and luminescence was measured using a luminometer with an integration time of 10s.

### qPCR analysis

As a spike-in, 10ng of firefly luciferase mRNA was added to isolated extracellular vesicles prior to RNA extraction. RNA was purified from isolated extracellular vesicles using TRIzol – LS (Thermo Fisher Scientific) following manufacturer’s guidelines. cDNA synthesis was carried out using random hexamers and Improm-II reverse transcriptase (Promega). The resulting cDNA was used for qPCR via iQ SYBR Green Supermix (Bio-Rad). The primers used for qPCR are as follows: Arc (AAGTGCCGAGCTGAGATGC, CGACCTGTGCAACCCTTTC); firefly luciferase (TTCGACCGGGACAAAACCAT, ATCTGGTTGCCGAAGATGGG). qPCR was done on CFX96 Touch Real-Time PCR Detection System (Bio-Rad).

### Cryo-EM specimen preparation, data-collection, and analysis

Extracellular vesicles were flash frozen and stored at -80 °C until use. Prior to preparing cryo-EM grids, the sample was diluted to 1.5 mg/ml as determined by a Bradford assay. C-flat grids (Copper, 1.2/1.3, Protochips) were prepared by glow-discharged for 30 sec in a PELCO Easiglow glow-discharge unit at 15 mA. We applied 3 μl of EV in PBS to the grid and incubated for 60 sec before vitrification using an FEI Vitrobot Mark IV (ThermoFisher). The grids were blotted for 3 sec using blotting force 3 at 4 °C and ∼90% humidity and plunged in liquid ethane.

Images were collected using a 626 Gatan cryo-holder on a TF20 microscope (FEI) equipped with a K2 Summit (Gatan) direct detector. We automated the collection of data using SerialEM (Mastronarde, 2005). 60 movies (50 frames/movie) of each sample were acquired over 6.8 sec with an exposure rate of 14.0 e^-^/pix/sec, yielding a total dose of 56 e^-^ /Å^2^, and a nominal defocus range from -1.0 μm to -3.0 μm. Images were gain corrected using the method described by (Afanasyev et al., 2015) and motion-corrected in cisTEM (Grant et al., 2018). We measured the length and width of EVs in 20 consecutively collected images per sample manually in tdisp (https://sourceforge.net/projects/tigris/) and calculated means and standard deviations of counted vesicles per image.

### FLIR imaging

Thermal changes in the hindpaw of vehicle- or CFA-injected were visualized using a FLIR T31030sc thermal imaging camera (FLIR instruments). Animals were placed in acrylic boxes with wire mesh floors and imaged 24 h post CFA injection. Image analysis and quantification were performed using the FLIR ResearchIR Max 4 software available at http://support.flir.com/rir4.

### Ultrasound imaging

Power Doppler ultrasound images of the hindpaw were acquired using a high-frequency ultrasound system (Vevo 3100, FUJIFILM VisualSonics Inc, Toronto, Canada) equipped with an MX550D linear array transducer operating at 32 MHz. Data were collected from different planes by precise movement of the transducer along the elevational direction (step size = 38 μm) using a specialized 3D motorized stage. Data were processed using the Vevo software module (FUJIFILM VisualSonics) to calculate paw volume and percent vascularity.

### Statistical analysis

*In vitro* data were collected from three independent cell cultures and are shown as means ± s.e.m. Behavioral testing data are shown as means ± s.e.m. of at least six animals per group. For behavior experiments, sample size was estimated as n = 6 using G*power for a power calculation with 80% power, expectations of 50% effect size, with α set to 0.05. Graph plotting and statistical analysis used GraphPad Prism Version 7.0 (GraphPad Software). Student’s t test was used to compare two independent groups. Statistical evaluation for three independent groups or more was performed by one-way or two-way analysis of variance (ANOVA), followed by post hoc Bonferroni, Dunnett, or Tukey test, and the a priori level of significance at 95% confidence level was considered at P < 0.05. Specific statistical tests used are described in figure legends. The significance of gene expression level changes before and after the treatment was calculated using a 2-tailed Student’s t-test assuming unequal variances.

## ACKNOWLEDGEMENTS

This work was supported by NIH grants R01NS065926 (TJP), R01NS098826 (TJP), and R01NS100788 (ZTC). The University of Texas STARS program (TJP) and the IASP Early Career Research Grant (PBI). J.S. received support from the Chan Zuckerberg Ben Barres Early Career Acceleration Award and an NIH Director’s Transformative R01 NS115716.

## AUTHOR CONTRIBUTIONS

Z.T.C., T.J.P., and P.B.I. conceived of the study. P.B.I., J.B.D.L.P., T.L., S.L., S.M., J.K.M., and N.K. performed experiments. A.W., P.R., and T.S., analyzed the sequencing data. Z.T.C., J.S., T.J.P., O.S., and P.B.I. wrote the manuscript. All authors approved the manuscript.

## SUPPLEMENTARY MATERIALS

